# Potential Benefits of Ketone Therapy as a Novel Immunometabolic Treatment for Schizophrenia

**DOI:** 10.1101/2024.05.23.595523

**Authors:** Karin Huizer, Shubham Soni, Mya A. Schmidt, Nuray Çakici, Lieuwe de Haan, Jason R. B. Dyck, Nico J. M. van Beveren

**Author notes:** Contributed equally (Jason R. B. Dyck, Nico J. M. van Beveren).

## Abstract

**Rationale:** Current treatment options for patients with schizophrenia-spectrum disorders (SSD) remain unsatisfactory, leaving patients with persistent negative and cognitive symptoms and metabolic side effects. Therapeutic ketosis was recently hypothesized to target the bio-energetic pathophysiology of SSD. However, neuro-inflammation plays an important role in the pathobiology of SSD as well. Ideally, novel treatments would target both the bio-energetic, and the inflammatory aspects of SSD. In this study, we aimed to investigate the effects of ketone bodies on neuro-inflammation in an acute inflammation mouse model.

**Methods:** 8-week-old male C57BL/6 N mice (n=11) were treated with either ketone ester (KE) or vehicle for 3 days. On day 3, a single intraperitoneal injection of lipopolysaccharide (LPS) or phosphate buffered saline (PBS) was administered. Mice were euthanized 24 h after LPS/PBS injection. Whole brain gene expression analysis using RT-PCR was done for *Tnf-a, Il-6* and *Il-1b*.

**Results:** LPS caused a potent transcriptional upregulation of *Tnf-a, Il-6* and *Il-1b* in the vehicle-treated mouse brain compared to PBS-injected controls. KE strongly and significantly attenuated the increased transcription of pro-inflammatory cytokines (*Tnf-a, Il-6* and *Il-1b*) in the brain upon LPS injection compared to vehicle.

**Conclusions:** KE potently dampened neuro-inflammation in this acute inflammation mouse model. Ketone therapy holds great promise as a treatment for SSD patients by simultaneously targeting two main pathophysiological disease pathways. We encourage more research into the immunometabolic potential of therapeutic ketosis in SSD.

**Highlights:**

- A brain bio-energetic deficit and neuro-inflammation are involved in schizophrenia
- Ketone therapy is being investigated as a bio-energetic treatment of schizophrenia
- Ketone ester inhibits neuro-inflammation in an acute inflammation mouse mode
- Therapeutic ketosis could target both pathophysiological pathways in SSD
- The Immunometabolic potential of ketone therapy for SSD warrants further attention

---

Approximately 1% of the world’s population is affected by schizophrenia-spectrum disorders (SSD) [1]. SSD is characterized by hallucinations, delusions, disorganized thought and behavior (positive symptoms), emotional blunting, lack of motivation and social withdrawal (negative symptoms) and cognitive impairments [2]. The latter two are largely responsible for the high morbidity associated with this illness. Current pharmacological treatment is aimed at modifying dopaminergic neurotransmission and tackles hallucinations and delusions with reasonable efficacy. However, even with current antipsychotic medications, negative and cognitive symptoms can persist, or are worsened [3]. Thus, novel treatments for negative and cognitive symptoms are needed to improve the prognosis of SSD patients.

Several biological models have been presented concerning the biological underpinnings of SSD. We here discuss the immune and neurometabolic models, both with implications for novel treatment strategies. First, the immune model describes that immune dysfunction resulting in disturbed neurodevelopment and neuro-inflammation in the pathophysiology of SSD. The immune model is supported by several converging lines of research [4-6]. In post-mortem brain tissue, elevated mRNA and protein levels of pro-inflammatory cytokines (esp. IL-6, TNF-α, IL-1β, and IFN-γ), as well as increased numbers of central nervous system-associated macrophages (CAMs) and activated microglia are found in at least a subset of SSD patients [6-10]. Pro-inflammatory cytokines (again esp. IL-6, TNF-α, IL-1β, and IFN-γ) are consistently found elevated in the blood [5, 11] and cerebrospinal fluid [12] of schizophrenia patients. Circulating pro-inflammatory cytokines are highest in treatment-naïve patients [13] and during acute disease phases, but their concentrations remain elevated in the chronic, treated disease phase compared to healthy controls [5]. Patients could therefore clinically benefit from further normalizing systemic and neuroinflammation.

Second, the neurometabolic model postulates that a bio-energetic deficit in the brain causes SSD, particularly the negative and cognitive symptoms [14, 15]. This model is supported by several findings of glucose metabolism disturbances systemically (including hyperglycemia, hyperinsulinemia, insulin resistance) [11], and in the brain (impaired glucose transport across the blood-brain barrier, disrupted metabolic astrocyte-neuron coupling, impaired glycolysis, deficits in the pentose-phosphate pathway, tricarboxylic acid cycle and oxidative phosphorylation [14]). The brain is heavily dependent on glucose both as an energy source, and as precursors to neurotransmitters [16]. Even subtle impairments in glucose metabolism will therefore negatively affect brain function [14, 16].

Since the immune system consists of metabolically highly active cells, there is a lot of functional cross-talk between energy metabolism and immune function [17, 18]. As such, both pathophysiological mechanisms are interrelated in SSD. While antipsychotic medications partially decrease pro-inflammatory cytokine levels [5, 13], most cause unwanted effects including insulin resistance, hyperglycemia, weight gain and metabolic syndrome [19]; these metabolic side effects further increase morbidity and reduce life expectancy in this already vulnerable patient group [20]. In line with the hypothesis that a bio-energetic deficit generates negative and cognitive symptoms, antipsychotics appear to further impair brain energy metabolism, and often worsen pre-existing negative and cognitive symptoms in schizophrenia patients [14].

Therefore, the ideal therapeutic intervention for SSD would simultaneously reduce (neuro)inflammation and restore the brain’s bio-energetic dysfunction without causing harmful metabolic side effects. Such a treatment approach could theoretically also improve negative and cognitive symptoms in SSD. Fortunately, a new type of treatment strategy, called ketone therapy, has emerged that may be able to address both of these issues.

The bio-energetic model has led Henkel et al. to suggest that ketosis may act as a therapeutic intervention in schizophrenia [14]. Ketone bodies (β-hydroxybutyrate, acetoacetate, acetone) are formed by the oxidation of fatty acids in the liver during fasting/starvation or upon restricting dietary carbohydrates (i.e. a ketogenic diet). While the brain relies entirely on glucose as a source of energy during normal feeding conditions, it switches to preferentially metabolizing ketones when ketone bodies are present at sufficient levels in the circulation [21, 22]. The preferential metabolization of ketones over glucose spares glucose for fully glucose-dependent processes like nucleotide and protein synthesis, including neurotransmitter formation in the brain [23, 24]. Ketone bodies can therefore be used as an alternative energy source to glucose by the brain [25]. In SSD, bypassing deficits in glucose metabolism by providing ketone bodies could restore the brain’s bio-energetic deficit. This theory is supported by some recent small-scale studies indicating positive effects of the ketogenic diet on schizophrenia symptoms [26, 27]. However, one major limitation of this approach is patient compliance as maintaining a strict ketogenic diet is quite challenging. Thus, exogenous ketone supplementation appears to be a more practical approach to achieving elevated circulating ketone levels [24].

In addition to their energetic advantages, ketone bodies may also be beneficial to SSD in other ways. For instance, Soni et al. recently showed that ketosis induced by ketone ester supplementation exerts strong anti-inflammatory effects in an LPS-based acute inflammation mouse model. In that study, (pre)treatment for 3 days with a ketone ester ((R)-β-hydroxybutyl (R)-β-hydroxybutyrate monoester) elevated circulating ketones and potently dampened the heightened inflammatory response in the heart, kidneys, liver and blood in LPS-injected mice [28]. Importantly, the levels of cytokines commonly found elevated in SSD (TNF-α, IL-6, IL-1β, IFN-γ) were significantly attenuated by ketone therapy in these mice [28].

Remarkably, inflammation in the lungs was unaffected, indicating organ-specific anti-inflammatory effects of ketone bodies. Since the brain has unique immunological features [29], it remains unknown if ketone therapy using ketone esters can protect against neuro-inflammation.

In this brief communication, we now provide additional data from the same LPS/inflammatory mouse model study by Soni et al. [28], focusing on the anti-inflammatory effects of ketone bodies in the brain. Whole brain (n=8-11/group) mRNA expression analysis was done for *Tnf-a, Il-6* and *Il-1b* according to methods described before [28]. These cytokines were selected because of their consistently reported elevation and involvement in SSD [5-7, 9-12]. As expected, LPS caused a potent transcriptional upregulation of *Tnf-a, Il-6* and *Il-1b* in the vehicle-treated mouse brain compared to PBS-injected controls (**Fig. 1**), confirming the robust neuro-inflammatory effects of LPS [30]. (Pre)treatment with ketone ester for 3 consecutive days strongly attenuated the increased transcription of pro-inflammatory cytokines (*Tnf-a, Il-6* and *Il-1b*) in the brain upon LPS injection (**Fig. 1**). Transcription of Tnf-a in whole brain was 10.4-fold lower (p=0.0017) in the ketone ester+LPS mice compared to vehicle+LPS mice, and Il-6 and Il-1b were 3.2-fold (p=0.0006) and 2.8-fold (p=0.0367) lower respectively.

**Fig. 1:**
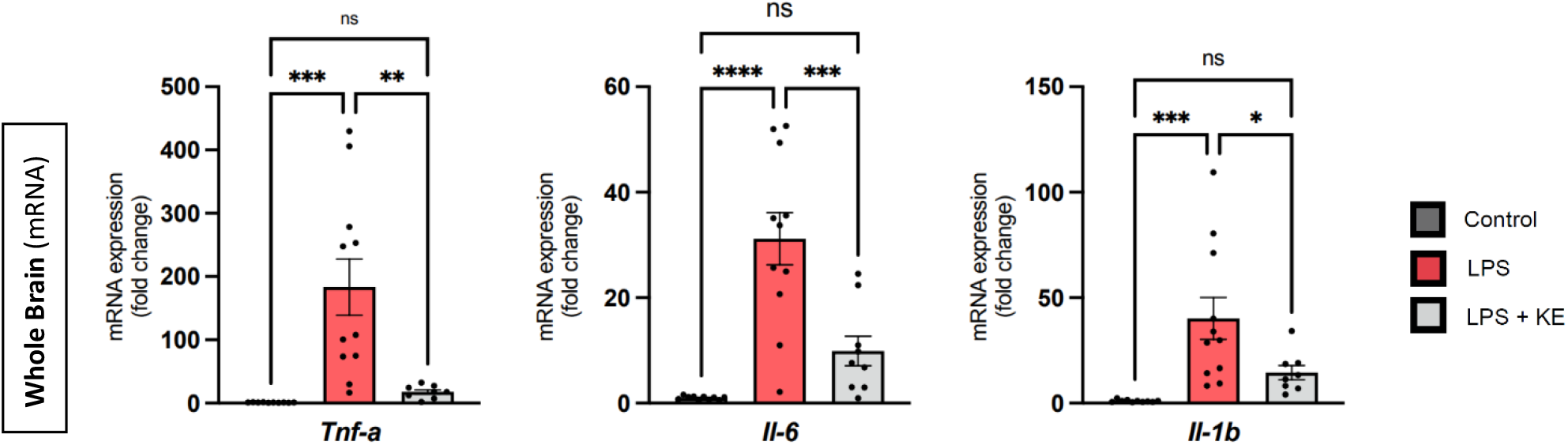
mRNA fold change of *Tnf-a, Il-6* and *Il-1b* in whole brain. LPS led to a strong upregulation of mRNA expression of brain *Tnf-a, Il-6* and *Il-1b*; the increase was largely prevented by prior KE supplementation. * p<0.05; ** p<0.01; *** p<0.001

Overall, these pre-clinical findings indicate that ketosis potently suppresses neuro-inflammation, by dampening the transcription of key pro-inflammatory cytokines involved in the pathophysiology of SSD. The precise anti-inflammatory mechanisms and effects of ketosis in the brain remain to be determined.

One hypothetical mechanism of action includes anti-neuroinflammatory effects as a result of glycolytic inhibition by ketone bodies [24]. Glycolysis is required for the activation of various pro-inflammatory immune cells [31], including microglia [18] and M1-macrophages [31] in the brain. Glycolytic inhibition by ketone bodies could therefore prevent LPS-induced proliferation/activation of microglia [30] and polarization of CAMs into the pro-inflammatory M1 subtype. Future studies using techniques allowing for high-plex expression analysis while preserving the spatial organization of the various immunological niches in the brain (e.g., the meningeal, choroid plexus, perivascular, parenchymal niches) [29] could help shed light on the anti-neuroinflammatory mechanism of action of ketosis.

Regardless of their precise mechanisms of action, increasing evidence is pointing towards the capacity of ketone bodies to simultaneously restore both the brain’s bio-energetic deficit in SSD, and reduce (neuro)inflammation, without causing harmful metabolic side effects. With current treatment options improving (neuro)inflammation, at the cost of further worsening pre-existing impairments in energy metabolism and overall metabolic health, ketone therapy therefore holds great promise as a treatment for SSD patients by simultaneously targeting two main pathophysiological disease pathways. If this promise holds true, therapeutic ketosis could help relieve the now so persistent and burdensome negative and cognitive symptoms in SSD patients, greatly improving their quality of life.

To investigate this premise, a pilot clinical trial on the use of exogenous ketone bodies in SSD patients will follow at Amsterdam University Medical Center upon receiving IRB approval, registered under NCT06426134 at ClinicalTrials.gov (https://clinicaltrials.gov/).

With this brief report, we hope to encourage more research into the immunometabolic potential of ketone bodies for the treatment of schizophrenia-spectrum disorders.

## Acknowledgments

JRBD receives funding from the Canadian Institutes of Health Research (CIHR), the Heart and Stroke Foundation of Canada, and Diabetes Canada. JRBD is a Canada Research Chair in Molecular Medicine. SS is supported by the CIHR Canada Graduate Doctoral Scholarship and the Izaak Walton Killam Memorial Scholarship. MAS is funded by the NSERC Canada Graduate Scholarships — Master’s program. NÇ is sponsored by an unrestricted personal grant to NJMvB.

